# Amyloid-β can activate JNK signalling via WNT-5A/ROR2 to reduce synapse formation in Alzheimer’s disease

**DOI:** 10.1101/2024.07.16.603668

**Authors:** Kevin Fang, Ehsan Pishva, Thomas Piers, Steffen Scholpp

## Abstract

Wnt signalling is an essential signalling system in neurogenesis, and recent studies have highlighted the critical role of this signalling network in regulating synaptic plasticity, neuronal survival, and neurogenesis, processes that are disrupted in Alzheimer’s disease (AD). From the Wnt network, the Wnt/β-catenin pathway has been studied for its neuroprotective role, and this is suppressed in AD. However, the involvement of the non-canonical pathway, which operates independently of β-catenin and involves the planar cell polarity (PCP), remains to be determined in AD.

In this work, we analyse the function of the orphan receptor tyrosine kinase ROR2, an essential co-receptor of the Wnt/PCP signalling pathway. We find that activation of WNT-5A/ROR2 signalling activates JNK signalling, reducing pre- and postsynaptic clusters on neurites in mature SH-SY5Y neurons. This observation is similar to SH-SY5Y neurons treated with the Amyloid-β peptide Aβ_1-42_ or DKK1, which are both increased in AD. Surprisingly, the effect of Aβ_1-42_ and DKK1 signalling on synaptogenesis can be mitigated by blocking ROR2 and JNK signalling, suggesting that Aβ and DKK1 signalling depends on ROR2/JNK signalling. Finally, we find an increase of WNT-5A/ROR2 clusters on neurites of iPSC-derived cortical neurons carrying the PSEN1 A75V mutation, known to enhance the pathological Aβ_42/40_ ratio. Simultaneously, the number of pre- and post-synaptic clusters decreased in the mutant line. Inhibition of ROR2/JNK signalling in PSEN1^A75V^ cortical neurons partially rescues the reduction in synaptogenesis, suggesting that ROR2 signalling may act in a positive feedback loop with Aβ_1-42_ and DKK1 signalling to augment JNK signalling as seen in AD.

## Introduction

Alzheimer’s disease (AD) represents one of the most prevalent neurodegenerative disorders, afflicting millions worldwide with a profound impact on memory, cognition, and overall quality of life^1^. AD is characterised by chronic neuroinflammation, synaptic dysfunction, loss of neurons, and brain atrophy. At the molecular level, the accumulation of amyloid-beta (Aβ) peptides and tau protein tangles in neurons are some hallmark characteristics of AD, contributing significantly to the disease’s pathogenesis^2–7^. However, the intricate mechanisms underlying synaptic dysfunction and subsequent neurodegeneration remain incompletely understood, hindering the development of effective therapeutics.

Within this complex disease landscape, the Wnt signalling network emerges as a critical regulator, offering insights into potential therapeutic targets for AD^8^. Recent studies have highlighted a crucial role of the Wnt/β-catenin signalling pathway in the stabilisation of synaptic connections, neuronal survival, and neurogenesis. In AD, these processes are disrupted, and indeed, Wnt/β-catenin signalling is down-regulated, and antagonists, such as DKK1 and 3, are upregulated^9–11^. Among the Wnt signalling pathways, the non-canonical Wnt/PCP signalling, particularly WNT-5A-mediated signalling, has gained emerging attention for its role in developing and maintaining the central nervous system (CNS)^12^. However, a link between dysfunctional Wnt/PCP signalling and AD is unclear.

WNT-5A, a ligand of the Wnt/PCP signalling pathway, regulates a wide range of cellular processes, including cell migration, polarity, and adhesion, through mechanisms distinct from the canonical Wnt/β-catenin pathway^13^. The function of WNT-5A is complex and involves interactions with various co-receptors, including the receptor tyrosine kinase-like orphan receptors ROR1/2. For example, WNT-5A/ROR2 signalling regulates the phosphorylation of Vangl2, which in turn facilitates the activation of the PCP/Jun N-terminal kinase (JNK) pathway. Interestingly, recent reports indicate that JNK signalling is upregulated in AD; has been linked to Aβ signalling^14, 15^; and JNK inhibitors have been shown to have a neuroprotective function and increase neuronal survival^16^. The Wnt signalling network and its regulating influence on the JNK pathway could play a role in modulating neuronal survival. However, the involvement of WNT-5A/ROR2 in regulating the effects of AD pathology and its interaction with Aβ signalling is less understood.

Given the intricate involvement of Wnt/PCP in neurogenesis and the potential impact of the JNK signalling pathway on AD progression, we observed differential expression of known Wnt regulators in association with AD amyloid pathology and identified WNT-5A/ROR/JNK. For our functional analysis, we used the human neuroblastoma cell line, SH-SY5Y treated with Aβ_1-42_ oligomers and compared the findings to iPSC-derived cortical neurons carrying the familial AD-linked PSEN1^A75V^ mutation, known to increase the plasma Aβ_42/40_ ratio, which is strongly linked to AD pathology^17–20^. In SH-SY5Y-derived neurons, we observed that increasing WNT-5A/ROR2 signalling leads to the activation of the JNK signalling pathway and enhanced sprouting of protrusions along neurites. Simultaneously, we also observe a reduction in the clustering of pre- and post-synaptic markers, suggesting a decrease in synaptogenesis. This phenotype resembles SH-SY5Y-derived neurons treated with Aβ_1-42_ peptides, together resembling some key aspects of AD. Surprisingly, inhibition of WNT-5A/ROR2/JNK signalling mitigates the phenotypes observed after Aβ_1-42_ and DKK1 treatment. In iPSC-derived cortical neurons carrying the PSEN1^A75V^ mutation, we found upregulation of WNT-5A/ROR2 clusters and enhanced JNK signalling. Supporting our results in the SH-SY5Y neurons, blockage of JNK signalling in iPSC-derived cortical neurons harbouring the PSEN1^A75V^ mutation can mitigate the loss of synaptic cluster formation. Our results suggest that in AD, an imbalance in the Wnt signalling network – here, an increased Wnt/PCP signalling - is linked to Aβ signalling in a positive feedback loop reinforcing JNK signalling.

## Results

### Expression of Wnt/PCP components is upregulated in an AD post-mortem brain

While the pathological hallmarks of AD include amyloid plaque deposition and Tau protein hyperphosphorylation, the underlying molecular mechanisms remain incompletely understood. A recent landmark study establishing a comprehensive single-cell atlas of the human prefrontal cortex across 427 individuals reported transcriptional differences associated with multiple measures of AD pathology and identified AD-associated alterations conserved across cortical layers and neuronal subtypes^21^ (Fig. 1A). We used this publicly available dataset to investigate the association of the Wnt signalling network in AD. Specifically, we focussed our analysis on the transcriptional profile of the excitatory cortical neurons of layers 3 to 4, which are affected before the onset of clinical symptoms^22, 23^. This neuronal population harbours crucial functions essential for memory and information processing and allows for a targeted exploration of Wnt signalling alterations potentially linked to early cognitive decline. In our analysis, we focussed on the differential expression of 114 genes from the Wnt network, focusing on ligands, receptors, and downstream effectors to elucidate the specific pathways potentially driving the disease process (Fig. 1A; Supplementary Table 1). Associated with increasing amyloid concentration and plaque burden (Fig. 1B, C), we observed an imbalance between Wnt/β-catenin signalling and Wnt/PCP/JNK signalling in the excitatory neurons in layers 3 and 4. For example, we found an upregulation of some critical components of the Wnt/PCP/JNK pathway, like the ligands WNT-5A/B, the cognate Wnt/PCP co-receptors of the ROR family and the downstream signalling factors JUN and ATF. Consistently, some canonical Wnt transcription factors were downregulated, like TCF7, TCF7L1, and LEF1. In parallel, known Wnt antagonists from the sFRP and DKK family were upregulated. As a positive control, we also found a significant upregulation of genes associated with Wnt and AD, such as CHD8 and members of the CSNK1 family. These findings suggest an increase of the non-canonical Wnt/PCP/JNK signalling pathway in excitatory cortical neurons in AD, potentially impacting neuronal health and function.

**Figure 1.**
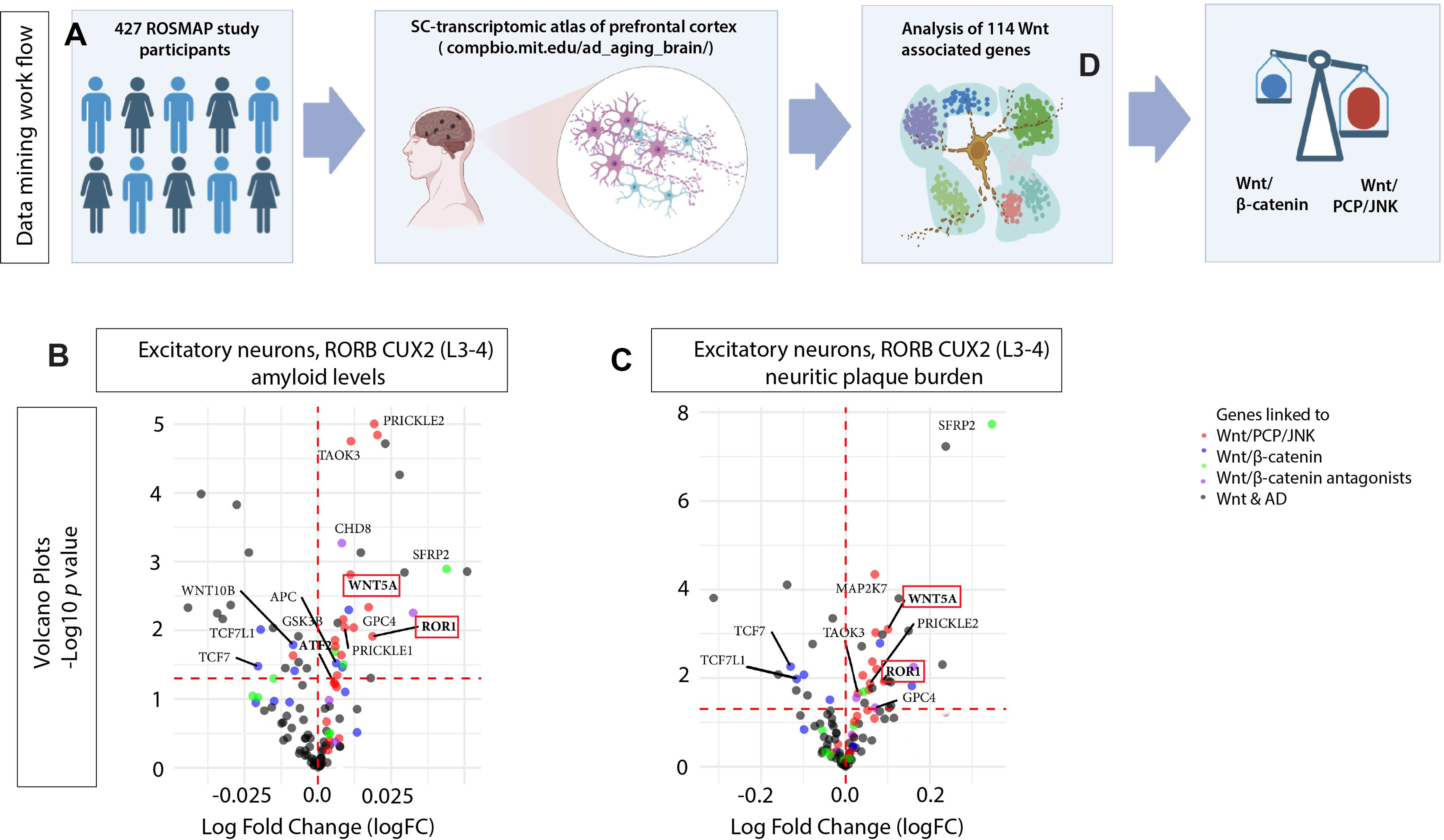
Differential gene expression study for Wnt-related genes in AD. (A) Cohort and snRNA-sequence profiling summary from 427 ROSMAP study participants along a spectrum of AD progression (based on Mathys et al., 2023, generated by Biorender). B, C, Volcano plots of the expression of 114 Wnt-related genes in excitatory neurons, RORB CUX2 of the layers 3 and 4 associated with amyloid levels (B), neuritic and diffuse plaque burden (C). The colour code indicates association with Wnt/PCP/JNK (red), Wnt/β-catenin (blue), Wnt/β-catenin antagonists (green), and AD-related genes (purple). The horizontal dashed line corresponds to *p* value < 0.05.

### Expression of WNT-5/ROR2 is enhanced in Aβ-treated SH-SY5Y cells

To determine such an effect of Wnt/PCP/JNK signalling in an AD context *in vitro*, we then established an SH-SY5Y-derived neuronal cell culture system. SH-SY5Y cells were plated on poly-D-lysine and laminin-coated culture dishes in serum-free media supplemented with nerve growth factor (NGF) and brain-derived neurotrophic factor (BDNF) to promote neurite outgrowth. After an initial proliferation period of 7 days, retinoic acid (RA) was introduced to the media to induce terminal differentiation. The extended differentiation period of up to 60 days allowed for the formation of extensive neuronal networks and expression of mature neuronal markers (Fig. 2A). After DIV 52, we found expression of Class III β-tubulin (TUJ-1) at high levels throughout the entire neuron, including dendrites, axons, and the cell body and Microtubule-associated protein 2 (MAP2) primarily localised in the dendrites and cell body of mature neurons. The differentiated and mature SH-SY5Y neurons were then used to determine the expression of the Wnt/PCP/JNK key regulators, WNT-5A/B and ROR2. In our IHC experiments, we found localisation of WNT-5A/B and ROR2 individually and in prominent clusters predominantly localised to crossing neurites (yellow arrows) on neurites (Fig. 2B, Supplementary Figure 2). To mimic an AD background chemically, we treated the neurons with 4 µM Aβ oligomers (Aβ_1-42_) or 200 ng/ml DKK1 protein for 24 h. We found that ROR2 mRNA expression, as well as ROR2 protein localisation on neurites, is significantly increased in SH-SY5Y neurons treated with AβO_1-42_ and DKK1 (Fig. 2D, E). The number of ROR2 mRNA transcripts increased approximately 1.60 and 1.53-fold, whereas the fold change for ROR2 protein puncta was 2.82 and 2.32, respectively.

**Figure 2.**
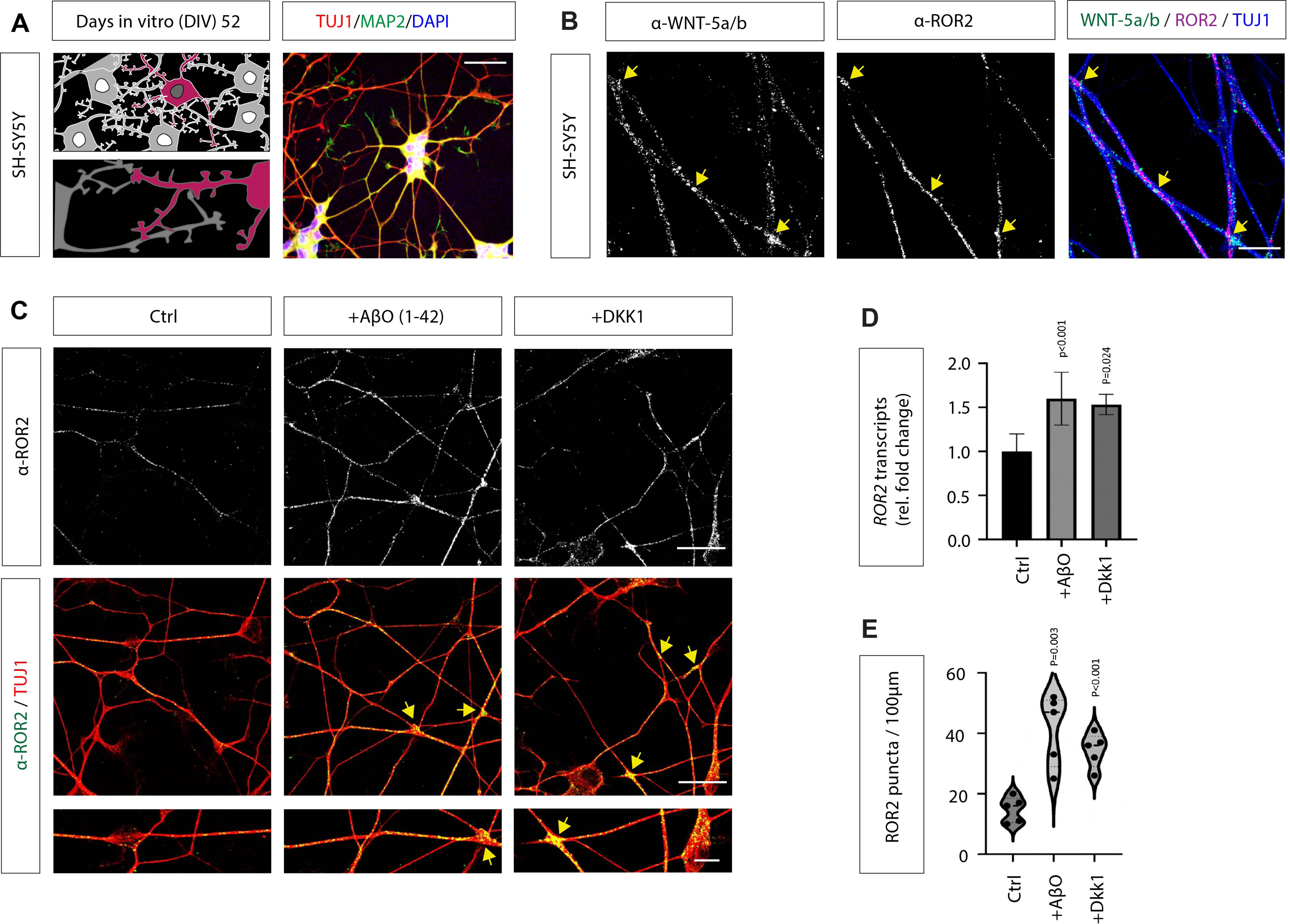
The cognate receptor ROR2 is co-localized with the Wnt ligands WNT-5A/B on dendrites of SH-SY5Y neuronal cells and is increased in an AD context. (A) SH-SY5Y cells were differentiated into neuron-like cells for 52 days and characterised by TUJ-1 (red), MAP2 (green) and DAPI (blue). Scale bar = 5 μm. (B) IHC co-staining of mature SH-SH5Y neurons to map the expression of ROR2 and WNT-5A/B. ROR2 is localised on neural dendrites and shows a robust co-localisation with WNT-5A/B (yellow arrows). Scale bar=5μm. (C) IHC shows endogenous ROR2 expression after treatment with indicated compounds in SH-SH5Y neurons. Quantification of Ror2 mRNA transcripts (D) and ROR2 protein puncta (E) demonstrates the upregulation of Ror2 after treatment with Aβ and DKK1. qRT-PCR was repeated 2 twice. 3-5 dendrites of each sample were selected to quantify ROR2 puncta and data from 3 biological replicates are displayed, including an unpaired Student-t-test with indicated *p* values.

### Soluble AβO can activate Wnt/PCP/JNK in a ROR2-dependent manner

WNT-5A and ROR2 are crucial components of the Wnt/PCP/JNK signalling pathway. Therefore, we tested if treating SH-SY5Y neurons with AβO_1-42_ and DKK1 influences JNK signalling. Thus, we used a JNK signalling reporter system to quantify the Wnt/PCP response in receiving neurons. Specifically, we used the JNK kinase translocation reporter, JNK KTR-mCherry, to monitor alteration in the JNK signalling cascade^24^. JNK-KTR-mCherry localises to the nucleus (N) in its dephosphorylated state (low JNK activity, Fig. 3A). Upon activation by phosphorylation, it shuttles to the cytoplasm (C) within minutes. To decipher the response of SH-SY5Y neurons to AβO_1-42_ and DKK1 transfected with the JNK-KTR-mCherry, the C: N ratio was calculated as an indicator of JNK signalling strength in near real-time^25^. SH-SY5Y neurons treated with AβO_1-42_ and DKK1 translocate the KTR into the cytoplasm, and the C: N ratio is significantly upregulated. AβO_1-42_ treated neurons showed a 3.02-fold increased C: N ratio, and DKK1-treated cells showed a 3.04-fold increase, indicating an activation of the JNK signalling pathway (Fig. 3B, C). As a positive control, we treated the neurons with WNT-5A protein, which led to a 2.65-fold increase, similar activation of the ROR2/JNK signalling indicated by the translocation of the KTR reporter.

**Figure 3.**
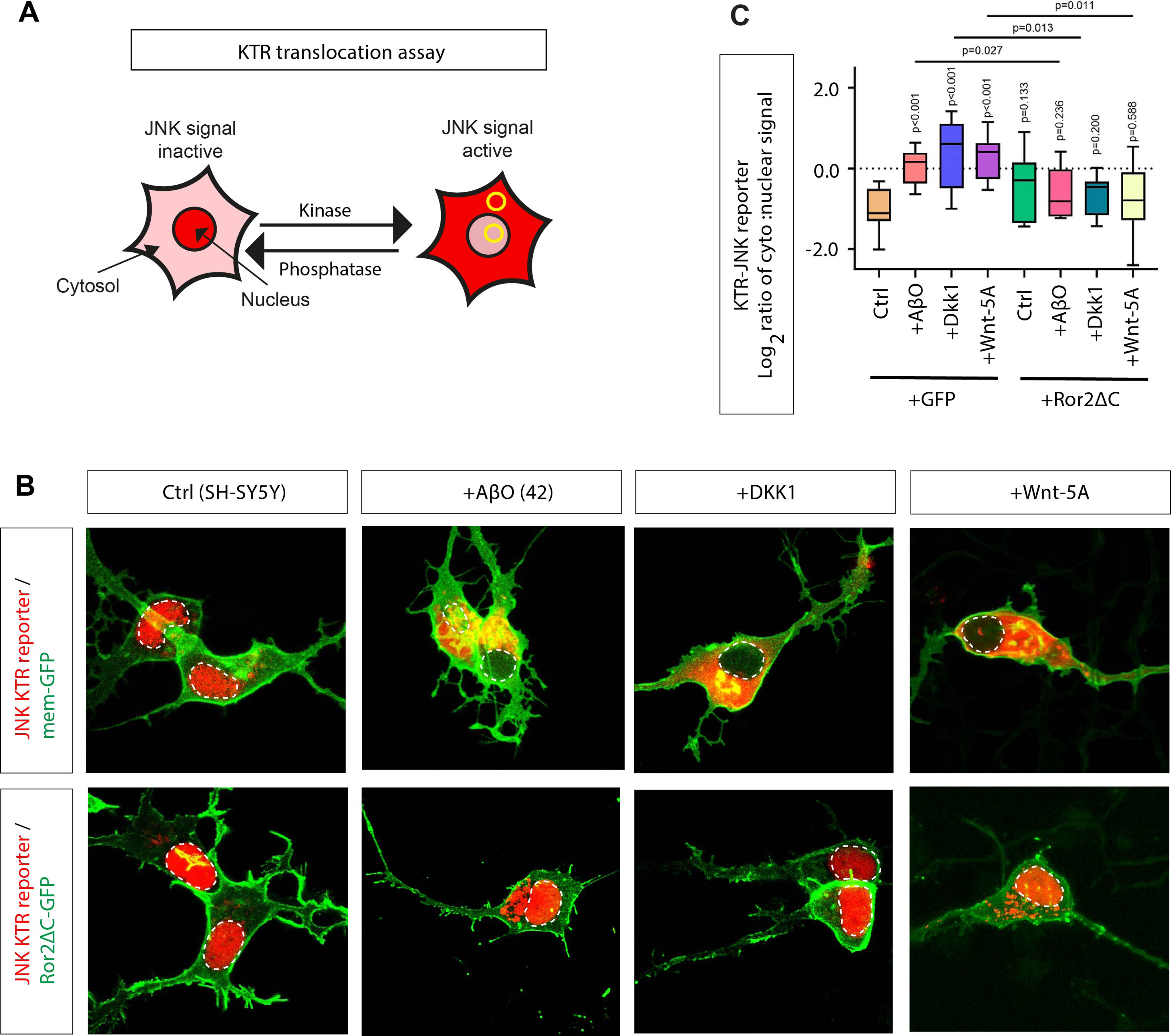
ROR2 facilitates JNK signalling in SH-SY5Y neurons in AD conditions. (A) Schematic representation of the functioning of the KTR-mCherry reporter. The reporter is localised to the nucleus at low JNK activity, while activated, phosphorylated JNK-KTR mCherry translocates to the cytoplasm. The signal ratio of cytoplasm: nucleus (C: N) was used to quantify JNK signalling activation. (B) Live images of SH-SY5Y neurons transfected with JNK-KTR-mCherry and Gap43-GFP or ROR2-ΔCRD. Soluble AβO_1-42_, DKK1 and WNT-5A protein were added at 24 h prior to imaging. The white circle indicates the nucleus. (C) The C: N ratio of 8 cells of 3 biological repeats were calculated, and an unpaired Student-T significance test with indicated *p* values is displayed. (C: N average values: ctrl: 0.49, +Αβ:0.98, +DKK1:1.00, +WΝΤ-5A:0.80, ROR2-ΔCRD:0.57, ROR2-ΔCRD+Αβ:0.59, ROR2-ΔCRD+DKK1:0.75, ROR2-ΔCRD+WΝΤ-5A:0.69.)

Next, we asked if the activation of the JNK signalling pathway depends on the Wnt/PCP receptor ROR2. Therefore, we co-transfected a dominant negative ROR2 construct lacking the intracellular kinase domain (ROR2-ΔC) together with the KTR reporter. We observed that neither AβO_1-42_, DKK1, nor WNT-5A can activate the JNK signalling significantly if the ROR2 function is blocked (Fig. 3B, C). These results suggest that, in an AD context, when AβO_1-42_ and DKK1 are upregulated, neurons can activate JNK signalling. However, activation of JNK signalling is facilitated by functional ROR2 receptors.

Next, we wanted to understand the consequences of upregulated Wnt/PCP/JNK signalling on synaptogenesis. Therefore, we treated SH-SY5Y neurons with AβO_1-42_ and DKK1 and investigated the expression of the presynaptic marker Synaptophysin-1 (Syp-1) and the postsynaptic marker Glutamate receptor ionotropic, AMPA1 (GluA1). We found that the number of clusters of Syp-1 and GluA1 on neurites is significantly reduced upon treatment with AβO_1-42_ and DKK1 (Fig. 4A, B). Similarly, we found a significant reduction of co-localised Syp-1/GluA1 clusters, suggesting decreased synaptogenesis. We then asked whether the observed phenotype depends on the activation of the Wnt/PCP/JNK pathway. To assess the role of JNK signalling in this process, we treated SH-SY5Y neurons with AβO_1-42_ or DKK1 and subsequently with a small molecule inhibitor of JNK kinase activity. SP600125 specifically inhibits all three kinases JNK1-3 (MAPK8-10) within minutes without inhibition of ERK1 or -2, phospho-p38, or ATF2^26^. Blockage of JNK signalling did not alter the number of co-localised Syp-1/GluA1 clusters in control neurons. However, if pre-treated with AβO_1-42_ and DKK1, SP600125 led to a significant increase in the emergence of Syp-1/GluA1 clusters in SH-SY5Y neurons (Fig. 4C, D). This data suggests that the down-regulation of Syp-1/GluA1 clusters in SH-SY5Y neurons by AβO_1-42_ and DKK1 depends on JNK signal activation.

**Figure 4.**
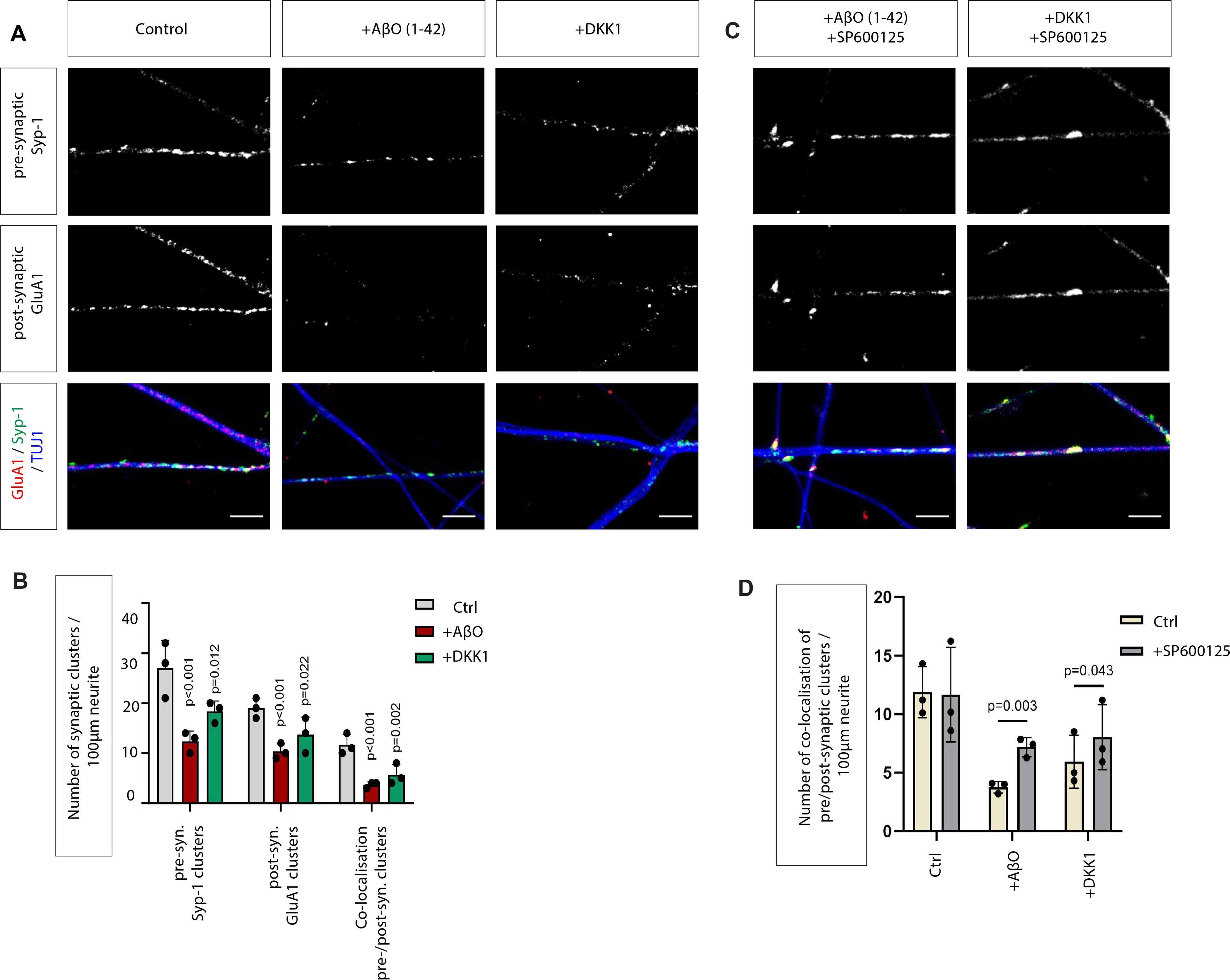
Treatment of SH-SY5Y neurons with AβO_1-42_ and DKK1 reduces synaptic marker localisation, which can be alleviated by blockage of JNK signalling. (A) Confocal images of SH-SY5Y neurons labelled with the presynaptic marker Synaptophysin-1(Syp-1, green), the postsynaptic glutamate receptor (GluA1, red) and β-tubulin (blue). Co-localization of Syp-1 with GluA1 indicates possible synaptic connections between SH-SY5Y neuron cells. Treatment with indicated compounds causes a loss of co-localisation. Scale bar = 5 μm. Quantification is shown in (B). (C) Confocal images show the clusters of synaptic markers pre-treated with JNK inhibitor SP600125. Quantification is shown in (D). (B, D) In the quantification, 3-5 dendrites of each sample were selected to quantify the number of puncta, and data from 3 biological replicates are displayed, including an unpaired Student-t-test with indicated *p* values.

### The Wnt/JNK receptor ROR2 expression depends on Presenilin 1 function

To complement the findings from the SH-SY5Y cells in a more pathophysiological context, we established iPSC-derived cortical neuron cultures (iNeurons). From the iNDI project, we obtained the parental KOLF2.1J iPSC line and genome-edited daughter lines carrying the familial Alzheimer’s Disease (fAD)-linked mutation PSEN-1^A75V27^. Comparing the isogenic cell lines provides a unique and powerful tool for dissecting the role of specific fAD-linked mutations known to increase the pathological AβO_42/40_ ratio. Here, we investigate the PSEN-1^A75V^ mutation, which increases the Aβ_42/40_ ratio and is associated with early-onset AD^28, 29^. After differentiation over 42 DIV, we found the expression of TUJ -1 and MAP2, indicating young cortical neuron phenotype (Fig. 5A). We used this neuronal cell culture system to compare the expression of ROR2 and WNT-5A between WT iNeurons and PSEN-1^A75V^ iNeurons. We observed that the number of ROR2 clusters is increased 2.18-fold in neurons carrying this PSEN-1 mutation, which consequently led to a 1.52-fold increase in WNT-5A/ROR2 clusters (Fig. 5B, C). These data suggest that an increase in the AβO_42/40_ ratio can induce an increase in ROR2 expression in human iNeurons. Interestingly, we found no alteration in WNT-5A clusters, suggesting that the production of the ligand is unaltered.

**Figure 5.**
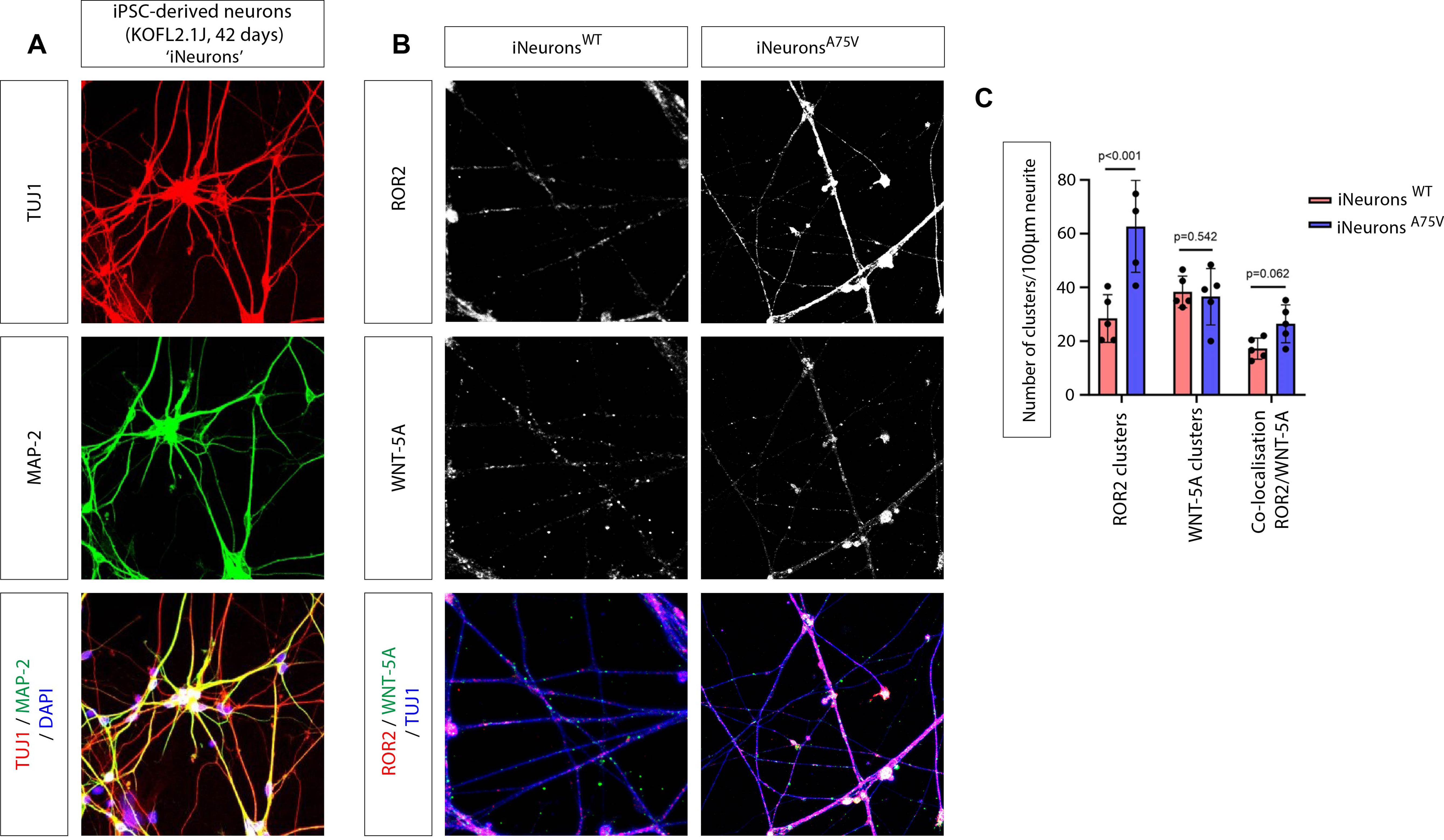
WNT-5-A/B / ROR2 expression in iPSC-derived cortical KOLF2.2 neurons. (A) iNeurons on 42 days show a robust expression of TUJ-1 (red), MAP2 (green) and DAPI (blue). (B) IHC staining of WT iNeurons and PSEN1^A75V^ iNuerons indicate the localisation of ROR2 and WNT-5-A/B. Scale bar = 5 µm. (C) In the quantification, 3-5 dendrites of each sample were selected to quantify puncta and 5 samples of 3 biological replicates are displayed, including an unpaired Student-t-test with indicated *p* values.

Next, we investigated the effect of AβO_1-42_ and DKK1 treatment on the clustering of pre- and postsynaptic markers in iNeurons. We found that treating iNeurons with AβO_1-42_ or DKK1 significantly reduced the formation of pre-synaptic Syp1 clusters by 1.61-fold and 1.42-fold, respectively (Fig. 6A, B). The post-synaptic GluA1 clusters were similarly decreased, here 1.60-fold and 1.39-fold, respectively. Consequently, the formation of co-localised Syp-1/GluA1 clusters was also reduced 1.70-fold and 1.36-fold (Fig. 6 A, B), in accordance with our previous observation in SH-SY5Y derived neurons. Together with the last experiment, these data suggest that AD-linked mutation PSEN-1^A75V^ in KOLF2.1J derived iNeurons may lead to an increase in ROR2 expression and, subsequently, a decrease of pre-and post-synaptic clusters.

**Figure 6.**
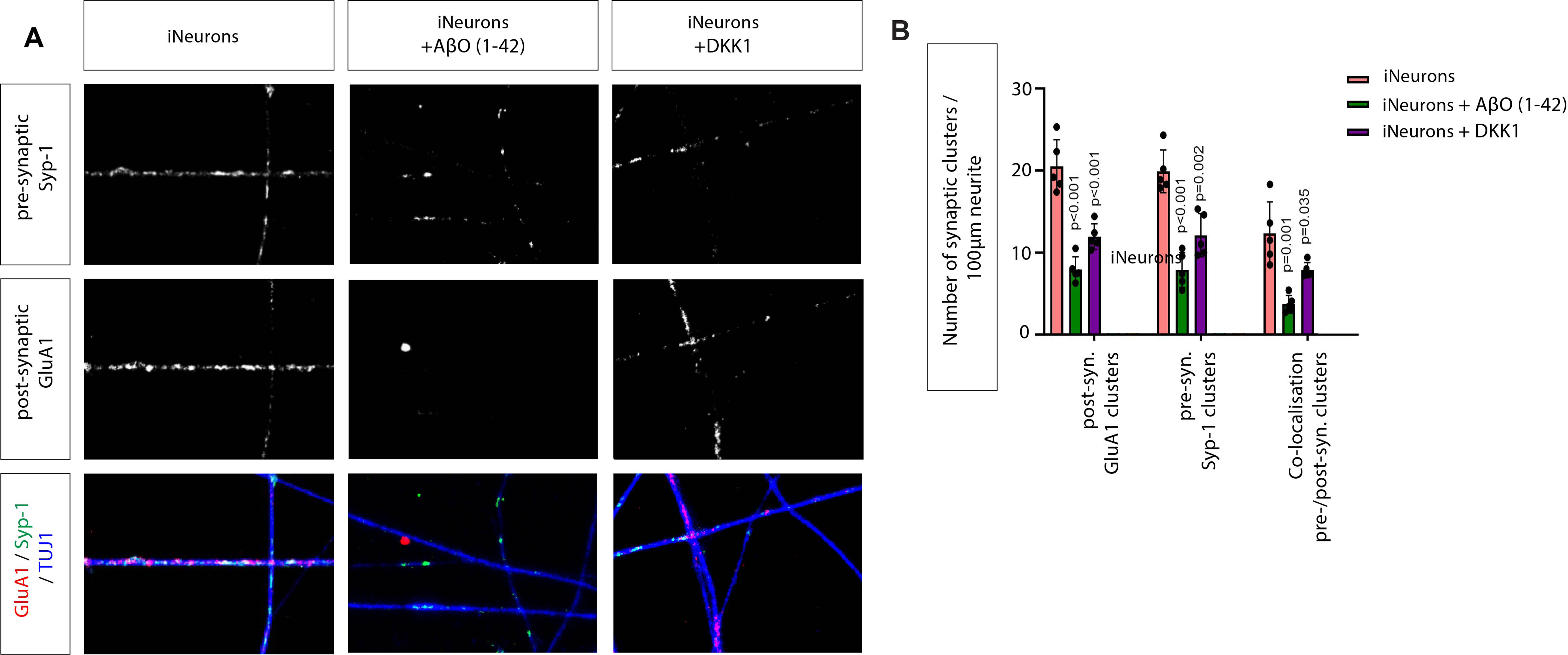
Treatment of soluble AβO_1-42_ and DKK1 affects synaptic clusters in iNeuons. (A) Confocal imaging of WT iNeurons IHC labelled with the presynaptic marker synaptophysin-1(Syp-1, green), the postsynaptic marker GluA1 (red) and β-tubulin (blue). Co-localization of Syp-1 with GluA1 shows potential synaptic connections. Scale bar = 5 μm. (B) In the quantification, 3-5 dendrites of each sample were selected to quantify puncta and 5 samples of 3 biological replicates are displayed, including an unpaired Student-t-test with indicated *p* values.

Based on our gene association analysis and data from the SH-SY5Y system, we hypothesise that Wnt/PCP/JNK signal activation reduces synaptic connections. Therefore, we treated WT iNeurons with WNT-5A protein and found that the number of pre-and post-synaptic clusters is significantly decreased, here by 1.38-fold (Fig. 7A, B). Finally, we wondered if PSEN1^A75V^ iNeurons show a reduced ability to cluster synaptic markers because of elevated Wnt/PCP/JNK signalling levels. Therefore, we treated the neurons with the JNK inhibitor SP600125 and found that we could partially rescue the pre- and postsynaptic puncta loss (Fig. 7A). We observed a significant 1.48-fold increase in puncta after blocking JNK signalling (Fig. 7B). Taking these findings together, these data suggest a central role for Wnt/PCP/JNK signalling in synaptic intergrity and an increase of Wnt/PCP/JNK in an AD context.

**Figure 7.**
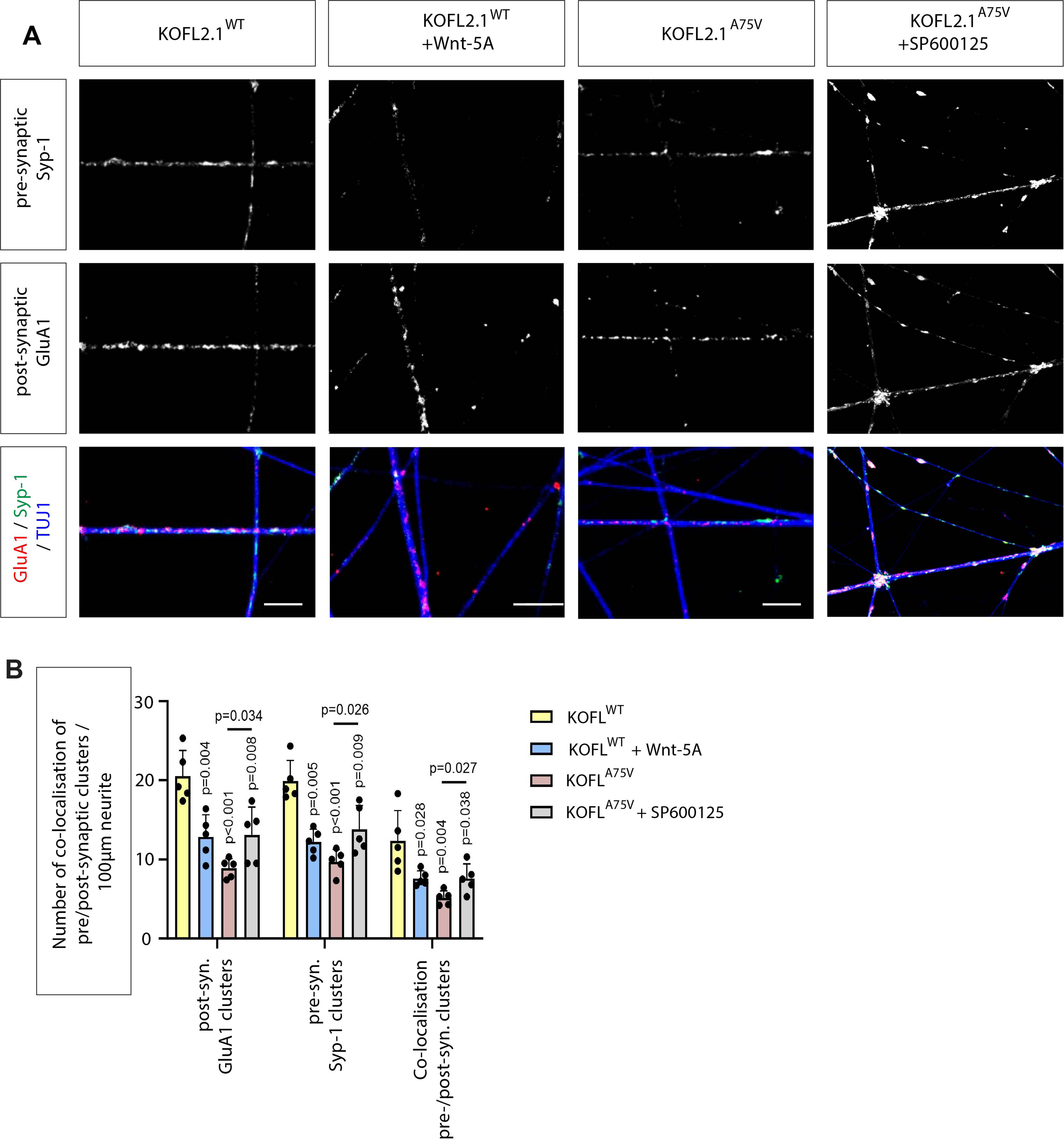
JNK inhibitor alleviates synaptic loss in PSEN-1^A75V^ iNeurons. (A) Confocal imaging of iNeurons labelled with the presynaptic marker synaptophysin-1(Syp-1, green), the postsynaptic marker GluA1 (red) and β-tubulin (blue). Co-localization of Syp-1 with GluA1 shows potential synaptic connections after treatment with the indicated compounds. Scale bar = 5 μm. (B) In the quantification, 3-5 dendrites of each sample were selected to quantify puncta and 5 samples of 3 biological replicates are displayed, including an unpaired Student-t-test with indicated *p* values.

## Discussion

AD is a progressive neurodegenerative disorder characterised by the accumulation of Aβ plaques and neurofibrillary Tau tangles, leading to neuronal loss and cognitive decline. In parallel, several major signalling pathways are misregulated during various stages of AD progression. For example, the canonical Wnt signalling pathway is suppressed in AD, whereas their cognate inhibitors, like DKK1-3, are massively upregulated^30^. These findings accelerated the research on the first specific Wnt inhibitors for Notum in AD^31^. Emerging evidence also highlights a critical role for the non-canonical Wnt signalling pathway in AD, which is regulated by WNT-5A. WNT-5A can initiate a cascade of events leading to the activation of the small GTPases RHO, RAC, and CDC42. In turn, this activates RHO Kinase (ROCK) and Jun N-terminal kinase (JNK). This cascade is crucial for various cellular processes in development and tissue homeostasis, including cell migration, differentiation, and apoptosis. However, how this signalling pathway is altered in AD and what the consequences are on neuronal function and health is unclear.

### An upregulation of Wnt/PCP/JNK is associated with AD

Here, we examined the association between expression of Wnt signalling network genes and AD at single-cell resolution^21^. This transcriptomic atlas of the aged human prefrontal cortex contains data from 427 ROSMAP study participants with varying degrees of Alzheimer’s disease progression, cognitive impairment, and AD pathology. Specifically, we analysed the expression for 114 Wnt/β-catenin and Wnt/PCP/JNK-related genes (Supplementary Table 1) in excitatory neurons around the cortical layers 3-4. We then selected the variables for plaque burden and amyloid levels and found an enrichment of genes related to the Wnt/PCP/JNK signalling system. In contrast, genes associated with the Wnt/β-catenin pathway are reduced.

In the subsequent analysis, we investigate the effect of Aβ signalling on young neurons, which complicates the direct translational ability of the findings. Based on the finding that WNT-5A/PCP/JNK is upregulated, we selected Wnt-5A, together with its cognate co-receptor ROR2, as a crucial signalling regulator for JNK signalling in AD for the subsequent analysis, as the ROR genes show redundancy in synaptogenesis^47^.

### ROR proteins are localised on neurites

To study their function in an AD context, we compared the effect of AβO in neuron-like cells differentiated from SH-SY5Y cells and iPSC-derived cortical neurons (iNeurons). While SH-SY5Y cells offer a readily available and relatively easy-to-maintain *in vitro* model, their limitations make iPSC-derived neurons a more robust and versatile tool for studying complex neuronal functions and diseases. Specifically, SH-SY5Y cells are a cancerous human neuroblastoma cell line that can exhibit an immature dopaminergic neuronal phenotype. Therefore, we complemented our analysis with iPSC-derived cortical neurons, which were genetically modified to introduce specific mutations, thus creating a more controlled model for studying disease mechanisms. Here, we established the KOLF2.1J iPSC line from the iNDI project^27^. These iNeurons are derived from a single genetic background, increasing the linkage of a specific mutation to a phenotype rather than inherent genetic variations between cell lines. This is particularly important in AD research, where genetic predisposition plays a significant role. Here, we used a cell line harbouring Familial Alzheimer’s Disease (FAD)-linked to the A75V mutation in PSEN-1, which is linked to early-onset AD and compared the effects to its isogenic control. Using these cell systems, we found that ROR2 expression is increased along neurites in an AD context.

Indeed, ROR2 is expressed in various neuronal populations, including in the hippocampus^32^. In cultured hippocampal murine neurons, ROR1 and ROR2 were detected during the neurite extension and synapse formation^33^. Both ROR proteins are initially localised on MAP2-positive dendrites and have been suggested to regulate their outgrowth. For example, murine hippocampal neurons, treated with antisense oligonucleotides or siRNAs targeting *ROR1* or *ROR2*, extended shorter minor processes with fewer branching points. However, inhibition of ROR1 in hippocampal neurons can reduce synaptogenesis^47^. The underlying mechanisms in regulating neurite extension in cortical neurons remain elusive.

Here, we show that the ROR2 receptor can be activated by WNT-5A/B in SH-SY5Y neurons and iPSC-derived human cortical neurons. Our data suggest that ROR2 can also be synergistically activated by high levels of AβO_1-42_ and DKK1, which have been observed in the AD context. Our results are further supported by the observation that KOLF2.1^A75V^-derived cortical neurons also significantly increase membranous ROR2 expression. Surprisingly, we found that inhibition of ROR2 can partially rescue the inhibitory effect of AβO_1-42_ and DKK1 signalling. These observations assign a key role for ROR2 in Aβ signalling and subsequently in AD.

### Wnt/PCP/JNK signalling in AD

Finally, our data suggest that WNT-5A/ROR2 are regulators of JNK signalling in the AD context. JNK signalling plays a significant role in regulating neuronal death, a hallmark of AD pathology. Previous studies have shown increased phosphorylated JNK (pJNK) expression in human postmortem brain samples from AD patients and a positive co-localization with Aβ^34^. Similarly, we observed upregulation of the expression of JNK-related genes by differential expression analysis (Fig. 1). Upon activation, JNK phosphorylates various substrates, including the transcription factor c-Jun, leading to the induction of pro-apoptotic genes^35^. The activation of JNK signalling has also been linked to the presence of AβO. It has been suggested that the interaction between AβO and neuronal receptors triggers a cascade of intracellular signals, culminating in the activation of JNK^36^. Specifically, JNK3 is strongly enriched in the nervous system and involved in neurite formation and neurodegeneration^37,38^. Phosphorylated JNK3 (p-JNK3) is highly correlated with AD pathology, such as promoting Aβ plaques and causing neuroinflammation, leading to nerve cell apoptosis. The level of JNK3 in the cerebrospinal fluid (CSF) of AD patients positively correlates with cognitive decline. A positive feedback loop regulates the formation of Aβ_1-42_: p-JNK phosphorylates APP at Thr668 to promote the production of Aβ_1-42,_ and Aβ_1-42_ reacts on JNK to promote its phosphorylation^39^. Finally, JNK signalling is involved in forming and aggregating neurofibrillary tangles (NFTs)^40^. Recent evidence suggests that inhibitors of JNK3 could be beneficial in AD^41^. Here, we show that WNT-5A/ROR2 is important for JNK signalling in cortical neurons. Blockage of ROR2 function or inhibition of JNK signalling can rescue the loss of pre- and postsynaptic marker localisation on neurites. In parallel, we found that increased levels of AβO also reduce pre- and postsynaptic marker localisation on neurites - again, this phenotype can be partially rescued by JNK inhibition. In parallel, AβO regulates GluA1 internalisation in HEK293T cells and primary cultured hippocampal and cerebrocortical neurons, suggesting an AβO-induced GluA1 suppression in neurons in early AD^42^. It is unclear if JNK signalling is also involved in this process.

In conclusion, we provide evidence of an interconnected signalling network suggesting a potential mechanism by which non-canonical Wnt signalling influences AD progression in the cortex. The modulation of JNK signalling by WNT-5A/ROR2, together with Aβ and DKK1, highlights the need to understand better the intertwined pathway interactions to develop targeted therapeutic strategies for AD.

## Acknowledgement

Research in the Scholpp lab is supported by the BBSRC (BB/S016295/1, BB/X008401/1, and an equipment grant, BB/R013764/1) and by the Living Systems Institute, University of Exeter. K.F. is supported by a Chinese Scholarship Council (CSC) studentship. E.P. is supported by ZonMw/Alzheimer Nederland grant (733050516). We would like to thank Corin Liddle (Bioimaging Centre Exeter) and Francesca Carlisle (Tissue Culture facility) for their technical support. We would further like to thank Akshay Bhinge (LSI Exeter) and Nicholas Nikolau (U Bath) for their critical comments on the manuscript.

## Material and Methods

### Plasmids and antibodies

The following plasmids were used in transfection experiments: pCS2+ Gap43-GFP (membrane-GFP); pCS2+-membrane-mCherry (Mattes et al., 2018); pCDNA-ROR2-GFP (Rogers et al, 2023), JNK KTR-mCherry (Regot et al., 2014), ROR2ΔC-GFP (Rogers et al, 2023).

The following primary antibodies were used for immunofluorescence: anti-ROR2 (Santa Cruz H-1), anti-WNT-5A/B (Proteintech, 55184-1-AP), anti-GluA1 (Synaptic Systems, 182 011), anti-Synaptophysin 1 (Synaptic Systems, 101 002), anti-β-3-tubulin (Synaptic Systems, 302 304), anti-MAP2 (Merck, MAB3418). The following AlexaFluor (Thermofisher) secondary antibodies were used for immunofluorescence: donkey anti-mouse 647(abcam; ab150107), donkey anti-rabbit 488 (abcam, ab150073), goat anti-Gpig 568 (abcam, ab175714), DAPI (Fisher, D1306).

### qPCR

RNA for qPCR was collected from cell pellets using the QIAGEN RNeasy kit according to the manufacturer’s instructions. qRT-PCR was then performed using the SensiFAST™ SYBR® Lo-ROX One-Step Kit with half volumes according to the manufacturer’s protocol and run using Applied Biosystems QuantStudio6 Flex. Primer sequences for ROR2 are: Forward 5’-ACTGGTCATCGCTTGCCTTT-3’, Reverse 5’-AGGCATGGAGACCTGTTTGT-3’.

### Cell culture

#### SH-SH5Y neuroblastoma cells maintenance

SH-SY5Y neuroblastoma cells were obtained from Akshay Bhinge’s lab. Cells were maintained in DMEM/F12 (Gibo), supplemented with 10% FBS. Lines were routinely passaged in a 1:5 split using 0.01% Trypsin (Thermofisher). The cell line was tested regularly for mycoplasma by endpoint PCR testing every 3 months and broth tests every 12 months.

#### Differentiation of SH-SY5Y neuroblastoma into mature neurons

Neurons were derived as previously described^43^. Briefly, SH-SY5Y neuroblastoma cells were plated onto 0.01% PDL and laminin-coated 6-well plates for starving culture by treating with DMEM/F12 plus 1% FBS supplemented with retinoic acid (10 μM). Media was changed every other day from D0-D8. On D8, cells were washed once by DMEM/F12 and cultures were replaced by neuronal differentiation media (DMEM/F12: Neurobasal in a 1:1 ratio, HEPES 10mM, N2 supplement 1%, B27 supplement 1%, Glutamax 1%) supplemented with retinoic acid (10 μM) and BDNF (10 ng/ml). On forwards D8, media was changed every other day and cells were maintained in culture until D40-60 for experimentation.

#### iPSC maintenance

iPSCs (KOLF2.1J) were obtained from the iNDI consortium (gift from Prof W. Skarnes). Lines were maintained as colonies on human ES-qualified Matrigel (Corning) in StemFlex (StemCell Technologies). Colonies were routinely passaged in a 1:6 split using EDTA and banked. All cell lines were tested regularly for mycoplasma by endpoint PCR testing every 3 months and broth tests every 12 months.

#### Differentiation of iPSCs into cortical neurons (iNeurons)

The differentiation process was described previously^44, 45^. Briefly, iPSCs were plated as colonies onto Matrigel and growing in neuronal differentiation media (DMEM/F12: Neurobasal in a 1:1 ratio, HEPES 10mM, N2 supplement 1%, B27 supplement 1%, Glutamax 1%, ascorbic acid 5uM, insulin 20ug/ml) supplemented with SB431542 (10uM) and LDN-193189 (0.2μM) from D0-D12, replacing media daily. On D12, cells were replated using accutase and growing in the same neuronal differentiation media supplemented with bFGF (20ng/ml), CHIR-99021 (1μM), and Y-27632 (50μM). On D13, media was changed to neuronal differentiation media supplemented with bFGF (20ng/ml), CHIR-99021 (1μM), and changed daily until D18. Cells can be banked and expanded neural progenitor cells (NPCs) on D18. For differentiating NPCs into cortical neurons, cells were maintained in differentiation media supplemented with L-Ascorbic acid (200μM), BNDF (20ng/ml), GDNF (10ng/ml), and Compound E (0.1μM) for 7 days before Compound E was removed. After D25, cultures were replenished every 3-4 days with a 50:50 media change. D40-60, cells were stained for cortical neuron lineage markers and used for experimentation.

### Neuronal transfection and treatment

Neuronal cultures were transfected with plasmid using a calcium-phosphate method as previously described^45, 46^. A plasmid/CaCl_2_ (12.4mM) master mix was prepared for each combination to be transfected in Hank’s balanced salt solution (HBSS). HBSS at the ratio of 1/8 of the DNA: CaCl_2_ mixture was added gradually to prevent the formation of large transfection complexes. The mixture was incubated for 15 minutes at RT, then added to cultures and incubated at 37°C for 4.5 h, before a sodium acetate (300mM in DMEM: F12) was added at 37°C for 5 minutes to dissolve precipitated complexes. Cultures were maintained in neuronal differentiation media supplemented for 24-48 h prior to imaging analyses.

### Pharmacological treatments of neuronal cultures

WNT-5A and DKK1 human recombinant protein was reconstituted at 100 μg/ml in PBS and 0.1% BSA (Biotechne). Amyloid β (1-42) peptide (Bachem, 4090148.05) was solubilised in DMSO at 1 mM, diluted to 100 mM in DMEM/F12, and allowed to oligomerise at RT for 16 hr. 200 ng/ml WNT-5A, 200 ng/ml DKK1 and 4 μM Aβ_1-42_ were added and incubated at 37 °C for 24 h before imaging or fixed for immunocytochemistry.

### Antibody staining and image acquisition

Cells were plated onto coverslips, and following indicated treatment/incubation, cells were immediately fixed using modified MEM-Fix (4% formaldehyde, 0.25–0.5% glutaraldehyde, 0.1 M Sorenson’s phosphate buffer, pH7.4) for 7 min at 4°C. Aldehydes were subsequently quenched by incubation with NaBH4 (0.1% w/v) for 7 minutes at RT. Further quenching was performed by 3 x 10 min washes in PBS-Glycine (0.2M). Cells were then blocked and permeabilised in permeabilisation solution (0.1% Triton-X-100, 5% donkey serum, 0.1 M glycine in 1×PBS) for 1 hr at RT. Primary antibodies were diluted in incubation buffer (0.1% Tween20, 5% serum in 1×PBS) and coverslips mounted on 40 μl spots overnight at 4°C in a humid environment. Coverslips were then washed with 1x PBS 3x for 5 min before mounting on 40 μl spots of secondary antibodies diluted in incubation buffer for 1 hr at RT. Coverslips were washed 3 times for 5 min with 1x PBS before mounting onto glass slides using ProLong Diamond Antifade mountant (Invitrogen) and left to dry overnight at 4°C before imaging.

### KTR-mCherry based JNK reporter assay and analysis

SH-SY5Y neuron cells were transfected with JNK reporter KTR-mCherry and either/or Gap43-GFP and ROR2ΔC-GFP. For analysis, mean grey values were recorded for 4× ROI in the nucleus and 4× ROI in the cytoplasm per cell and an average was taken. The ratio of cytoplasmic/nuclear signal was then recorded.

### Image acquisition and statistical analyses

Confocal microscopy images were performed on an inverted Leica TCS SP8 X laser-scanning microscope using a 63x water objective and analysed using ImageJ-FIJI. For analysis of dendritic puncta, 3-5 neuronal dendrites within an ROI were selected randomly. The number of puncta was counted using the cell counter plugin in FIJI, and an average number was taken. The number of co-localised puncta was quantified using the synapcountJ plugin in FIJI. Protrusion length was measured from the tip of the filopodia to the base, where it contacted the neurite. In the case of branching protrusions, one branch (the longest) would be measured. All experiments/conditions were repeated in biological triplicates. Statistical analyses were performed using GraphPad Prism version 10.1.2. Comparisons between the two groups were analysed using the unpaired two-sided Student’s T-test.

### Post-mortem brain differential expression data acquisition

The association between Wnt pathway genes and overall amyloid levels, neuritic plaque burden, and diffuse plaque burden for excitatory neurons, specifically the RORB CLUX2 cell cluster (cortical layers 3-4), was obtained from the Mathys Lab GitHub repository (https://github.com/mathyslab7/ROSMAP_snRNAseq_PFC/tree/main/Results/Differential_gene_expression_analysis/). These results were generated using 427 AD and non-demented post-mortem PFC samples. The details of the single-cell RNA sequencing experiment and the differential expression analysis are described in Mathys et al. The Volcano plots were created using the ggplot2 R package (version 3.4.2).

